# Expression of the Phosphatase Ppef2 Controls Survival and Function of CD8^+^ Dendritic Cells

**DOI:** 10.1101/374652

**Authors:** Markus Zwick, Thomas Ulas, Yi-Li Cho, Christine Ried, Leonie Grosse, Charlotte Simon, Caroline Bernhard, Dirk H. Busch, Joachim L. Schultze, Veit R. Buchholz, Susanne Stutte, Thomas Brocker

## Abstract

Apoptotic cell death of Dendritic cells (DCs) is critical for immune homeostasis. Although intrinsic mechanisms controlling DC death have not been fully characterized up to now, experimentally enforced inhibition of DC-death causes various autoimmune diseases in model systems. We have generated mice deficient for *Protein Phosphatase with EF-Hands 2* (Ppef2), which is selectively expressed in CD8^+^ DCs, but not in other related DC subtypes such as tissue CD103^+^ DCs. Ppef2 is down-regulated rapidly upon maturation of DCs by toll-like receptor stimuli, but not upon triggering of CD40. Ppef2-deficient CD8^+^ DCs accumulate the pro-apoptotic *Bcl-2-like protein 11* (Bim) and show increased apoptosis and reduced competitve repopulation capacities. Furthermore, Ppef2^−/−^CD8^+^ DCs have strongly diminished antigen presentation capacities *in vivo*, as CD8^+^ T cells primed by Ppef2^−/−^ CD8^+^ DCs undergo reduced expansion. In conclusion, our data suggests that Ppef2 is crucial to support survival of immature CD8^+^ DCs, while Ppef2 down-regulation during DC-maturation limits T cell responses.

## Introduction

DCs are major antigen-presenting cells (APC) located in tissues and lymphoid organs, where they integrate environmental signals to initiate either immunity or tolerance. They express receptors for pathogen associated molecular patterns allowing early recognition of microbial intruders. By conveying this information to T lymphocytes, DCs are linking innate and adaptive immunity.

Conventional DCs (cDC) develop from committed DC precursors (pre-DC) in the bone marrow (BM), which seed tissues and develop into either CD8^+^/CD103^+^ DCs or CD11b^+^ DCs (Liu, Victora et al., 2009, Merad, Sathe et al., 2013). Lymphoid organ resident DCs consist mainly of CD8^+^CD11b^−^ and CD8^−^CD11b^+^ cells, while migratory DCs express CD103 and/or CD11b. Both, CD8^+^ DCs and CD103^+^ DCs are superior to present antigen via MHCI to CD8^+^ T cells (Bedoui, Whitney et al., 2009, Dudziak, Kamphorst et al., 2007, Hildner, Edelson et al., 2008), while CD11b^+^ DCs are better equipped to present antigen via MHCII to CD4 T cells (Dudziak et al., 2007). In addition, CD8^+^ and CD103^+^ DCs are able to efficiently load exogenous antigen onto MHCI for cross-presentation to CD8^+^ T cells (den Haan, Lehar et al., 2000). More recently it was shown that CD8^+^ DCs have the specific and non-redundant role to optimize CD8^+^ T cell responses during a later step in T cell priming (Eickhoff, Brewitz et al., 2015). Several mutant mice have been generated to study the functions of CD8^+^/CD103^+^ DCs or CD11 b^+^ DCs *in vivo* (Merad et al., 2013, Murphy, Grajales-Reyes et al., 2016). However, mice with mutations that affect either CD8^+^ or CD103^+^ DCs, but not both subtypes do not yet exist.

The maintenance of DC populations relies on constant replenishment by blood-borne precursors (Liu, Waskow et al., 2007, Waskow, Liu et al., 2008) and *in situ* cell division with 5% of lymphoid organ resident DCs undergoing cell division at any given time (Kabashima, Banks et al., 2005). The importance of tightly controlled DC-numbers becomes obvious when the system is disturbed artificially. Inhibition of DC apoptosis by interfering either with caspases (Chen, Wang et al., 2006), pro-apoptotic Bim (Chen, Huang et al., 2007) or cell-death inducing Fas (Stranges, Watson et al., 2007) *in vivo* caused DC-accumulation and autoimmunity. Similarly, artificial prolongation of DC-lifespans by Akt mutants (Park, Lapteva et al., 2006) or overexpression of anti-apoptotic Bcl-2 (Nopora & Brocker, 2002) enhanced immunogenicity of DCs. However, mechanisms naturally regulating the DC lifespan are less well described. DC-maturation by lipopolysaccharide (LPS) induces apoptosis by CD14-mediated NFAT activation (Zanoni, Ostuni et al., 2009) and down-regulation of Bcl-2 (Hou & Van Parijs, 2004). Also killing of DCs by primed cytotoxic T cells (CTL) has been described (Hermans, Ritchie et al., 2000), a mechanism which was observed for both, CD103^+^ and CD11b^+^DC (Daniels, Hyde et al., 2016). Ligands of the tumor necrosis factor superfamily bind to CD40 (Riol-Blanco, Delgado-Martin et al., 2009) or TRANCE (Wong, Josien et al., 1997) on DCs to prolong their survival (Kushwah & Hu, 2010). However, to the best of our knowledge other intrinsic DC life-cycle regulatory mechanisms are not known.

Previously, we characterized the promoter regions of CD11 c and DC-STAMP, two DC-specific markers, and identified an evolutionary conserved promoter framework, which also controls expression of Ppef2 (Edelmann, Nelson et al., 2011). Ppef2 is a poorly characterised phosphatase with three EF-hands typical for calcium-binding proteins and an IQ motif (Montini, Rugarli et al., 1997). In mice, Ppef2 is strongly expressed in the retina, but Ppef2 deficiency did not cause retinal degeneration (Ramulu, Kennedy et al., 2001), while Ppef2-orthologues prevent retinal degeneration in *Drosophila* (Steele & O’Tousa, 1990). Besides the Ca^2+^-binding of Ppef2 (rdgC) in *C. elegans* (Ramulu & Nathans, 2001) or the Calmodulin-binding of Ppef2 in human cells (Kutuzov, Solov’eva et al., 2002), it has been speculated that Ppef2 would be involved in stress-protective responses and could possibly positively regulate cell survival, growth, proliferation and oncogenesis as a “survival-phosphatase” (Andreeva & Kutuzov, 2009).

Here, we show that in the hematopoietic system of mice Ppef2 expression is confined to CD8^+^ DCs, but not tissue resident CD103^+^ DCs or other cells. Ppef2 is down-regulated rapidly after DC-activation with toll-like receptor (TLR) ligands, while DC-maturation via CD40 did not alter Ppef2-levels. We generated Ppef2-deficient mice and show that CD8^+^ DCs display increased apoptosis *in vitro* and *in vivo*. As a consequence, the Ppef2^−/−^ DCs displayed strongly reduced cross-presenting capacities *in vivo*. Our data identify Ppef2 as a molecular regulator for survival and function of CD8^+^ DC.

## Results

### Ppef2 expression is restricted to immature CD8^+^ DCs

To analyse Ppef2-expression and regulation in greater detail, we purified lymphocyte populations from the spleen and blood of mice for gene expression analyses. Ppef2-expression was restricted nearly exclusively to MHCII^+^CD11c^+^CD8^+^ DCs in the spleen (Fig. 1a), confirming earlier results (Edelmann et al., 2011). Other cells such as MHCII^+^CD11c^+^CD11b^+^ Esam^hi^ DCs (Lewis, Caton et al., 2011); Suppl. Fig. 1a), B cells and various distinct blood monocyte populations expressed only very low levels of Ppef2 mRNA (Fig. 1a). Ppef2-expression was not detectable in pDCs, CD4^+^ T cells and CD8^+^ T cells (Fig. 1a). Taken together, our analyses confirmed a highly specific expression of Ppef2 in CD11b^−^CD8^+^ DCs as suggested by *Immgen* expression data collection (Heng & Painter, 2008) and our own previous findings (Edelmann et al., 2011). Also GMCSF-cultured BMDCs expressed Ppef2 (Fig. 1b). Here, CD11c^+^ DCs of different maturation stages as characterised by CD86 and MHCII surface expression, showed an inverted relation between Ppef2-mRNA expression and DC-maturation (Fig. 1b). In these cultures, MHCII^lo^CD86^lo^ immature DCs expressed the highest amount, while MHCII^int^CD86^lo^ semi-mature DCs expressed lower levels and most mature MHCII^hi^CD86^hi^ DCs expressed lowest levels of Ppef2 mRNA (Fig. 1b). This Ppef2-expression pattern in spontaneously matured cultured BMDCs suggested that Ppef2 is down-regulated with increasing DC maturation. Also, GM-CSF- or Flt3L-cultured BMDCs, as well as CD8^+^ DC from spleens, stimulated with various toll-like receptor (TLR) ligands, such as LPS (TLR4), flagellin (TLR5), Poly(I:C) (TLR3), Pam3CSK4 (TLR1/2) or CLO97(TLR7/8), all downregulated Ppef2 mRNA (Fig. 1c). In marked contrast, BMDC-maturation by crosslinking of surface CD40 with anti-CD40 mAb alone (Fig. 1c) did not cause down-regulation of Ppef2-expression (Fig. 1c). Addition of LPS to anti-CD40 stimulated BMDCs caused down-regulation of Ppef2-mRNA, indicating a dominant effect of TLR-over CD40-signalling (Fig. 1c). Taken together, our data indicates that CD8^+^ DCs selectively express Ppef2 mRNA, which is down-regulated rapidly upon DC-maturation by TLR-signalling, but not by CD40-crosslinking.

**Figure 1.**
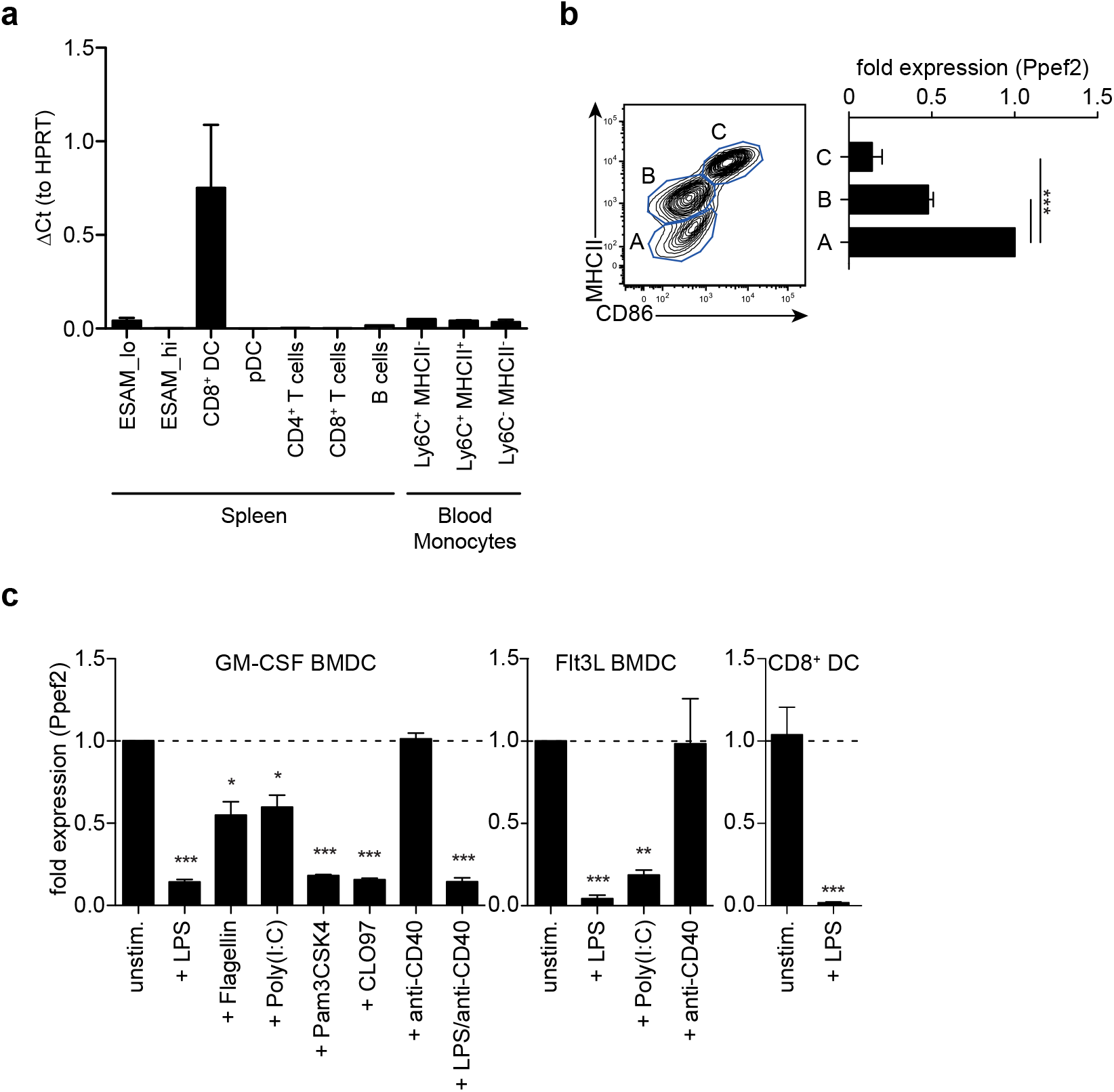
Ppef2 is predominantly expressed by CD8^+^ DCs. **(a)** Gene expression profiling of spleen and blood cells after sorting. Shown is the quantitative real-time PCR result for Ppef2 of three independent sort experiments (n=3 pooled mice per sort) ± SEM analyzed by the ΔCt method with HPRT as housekeeping gene. Cells were identified and sorted as ESAM^hi^ (CD11c^+^MHCII^+^CD11b^+^ESAM^hi^); ESAM^lo^ (CD11 c^+^MHCII^+^CD11b^+^ESAM^lo^); CD8^+^DC (CD11c^+^MHCII^+^CD11 b^−^CD8^+^); pDC (CD11b^−^ SiglecH^+^); CD4^+^ T cells (CD3^+^CD4^+^); CD8^+^ T cells (CD3^+^CD8^+^); B cells (CD19^+^B220^+^); blood monocytes NK1.1^−^B200^−^CD115^+^CD11b^+^Ly6G^−^ cells with differential Ly6C and MHCII expression as indicated. Gating strategies are shown in Supplemental Figure 1b. Ppef2 expression in GM-CSF cultured BMDCs of C57BL/6 mice after cell sorting on day 7 of culture based on the expression of CD11c, MHCII, and CD86. Error bars represent SEM of 3 independent experiments. **(c)** Ppef2 expression 16h after *in vitro* stimulation with the indicated TLR-ligands of GM-CSF- or Flt3L-cultured BMDCs of C57BL/6 mice, as well as spleen CD8^+^ DCs 16h after intravenous injection of 10μg LPS as determined by qPCR. Bar graphs with SEM represent pooled data from independently performed cell cultures (GM-CSF BMDCs: unstim., LPS (n=5); Flagellin, Poly(I:C), Pam3CSK4, CLO97, (*n*=4); anti-CD40, LPS^+^anti-CD40 (n=3); Flt3L BMDCs (*n*=3); sorted CD8^+^ DCs (*n*=4)). Statistical analysis was performed using Student’s t-test, with *p<0.05; **p<0.01; ***p<0.001.

### Generation of Ppef2-deficient mice

To explore the role of Ppef2 *in vivo*, we next generated Ppef2-deficient (Ppef2^−/−^) mice using the knock-out first strategy (Skarnes, Rosen et al., 2011) by introducing a gene-trap between exons 4 and 5 as well as flanking loxP-sites of exon 5 (Fig. 2a). DCs from Ppef2^−/−^ animals lacked Ppef-mRNA, as detected by quantitative PCR (Fig. 2b) as well as the analysis of reads from mRNA-sequencing of CD8^+^ DCs (Fig. 2c). Similar results were obtained from total splenocytes, where also no Ppef2-mRNA was detectable (data not shown). These analyses revealed complete lack of Ppef2-transcripts beyond exon 3, indicating successful knockdown of gene expression (Fig. 2b, c). Due to lack of functioning Ppef2-specific antibodies, further direct studies of Ppef2-protein expression were not possible. However, the introduction of a gene trap/lacZ construct allowed indirect monitoring of expression and regulation of Ppef2 using ß-galactosidase (ß-Gal) as a surrogate. FACS-analysis of various cell types from spleen, lymph nodes and thymus confirmed the highly selective expression of Ppef2/ß-Gal in CD8^+^ DCs of spleen and lymph nodes as well as CD8^+^ and CD11b^+^ DCs of thymus (Fig. 2d, e). Other cells such as CD8^+^ T cells, LN CD103^+^ and CD11b^+^ DCs, spleen CD11b^+^ESAM^hi^ and CD11b^+^ESAM^lo^ DCs, monocytes (Fig. 2d, e), CD4^+^ T cells, B cells and neutrophils (data not shown), did not express Ppef2/ß-Gal *in vivo*. Furthermore, lineage^−^MHCII^−^ CD11c^low^CD43^+^sirp-α^+^ DC-precursors (Suppl. Fig 1e)(Naik, D. et al., 2006), which give rise to both, CD8^+^ DCs and CD103^+^ DCs in lymphoid and non-lymphoid tissues (Liu et al., 2009), did not express Ppef2/ß-Gal (Fig. 2d, e). Interestingly, while CD8^+^ DCs were Ppef2/ß-Gal-positive, CD103^+^ DCs in lung and intestinal lamina propria tissue, which develop from the same pre-DC were Ppef2/ß-Gal-negative (Fig. 2d, e). This indicates that Ppef2/ß-Gal-expression is regulated after the pre-DC stage and might be sensitive to signals from tissue milieu or the environment. CD8^+^ DCs also showed strong down-regulation of Ppef2/ß-Gal upon stimulation with TLR4-ligand LPS *in vivo* (Fig. 2f), confirming the shut-down of Ppef2 expression upon maturation of DCs *in vivo*. These data demonstrate the successful knockout of Ppef2, its specific expression in the CD8^+^ DC subset and Ppef2 down-regulation upon DC-maturation.

**Figure 2.**
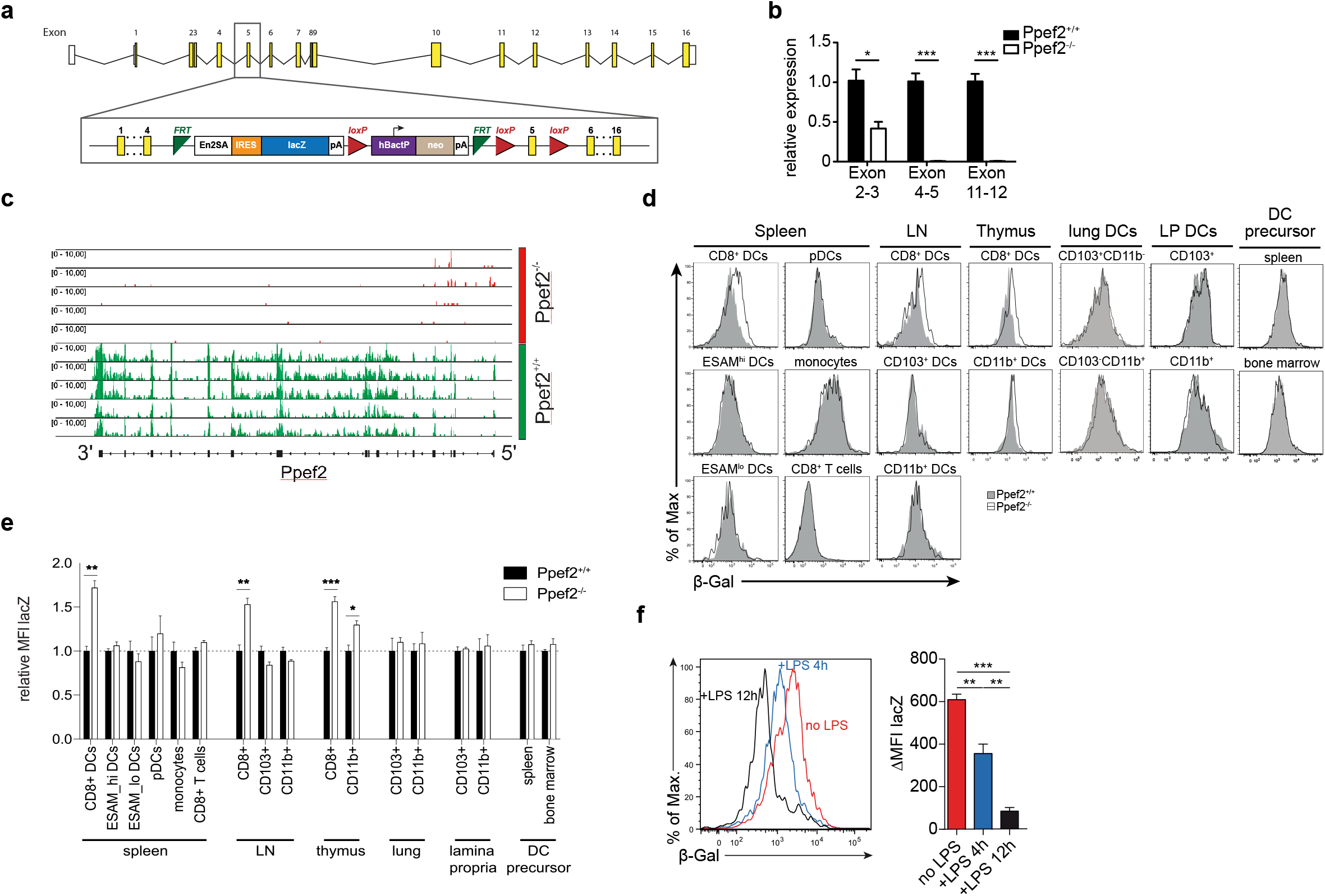
Ppef2-knockout strategy and LacZ-Reporter. **(a)** The Ppef2 locus with the knockout-first construct containing the gene trap between exon 4 and 5, as well as the floxed exon 5. **(b)** Ppef2 expression analysis of CD11c enriched splenocytes in Ppef2^+/+^ and Ppef2^−/−^ mice by quantitative real-time PCR. Three sets of intron-spanning primer pairs were used to amplify fragments from exons 2 to 3, 4 to 5 and 11 to 12. The ΔCt method was used to calculate the fold expression compared to Ppef2^+/+^ control samples. Data was normalized to HPRT (n=3 mice). **(c)** Genomic distribution of reads across the Ppef2 gene locus in Ppef2^+/+^ and Ppef2^−/−^ CD8^+^ DCs. **(d)** Measurement of ß-Gal activity by flow cytometry in different cell types of the spleen, lymph node, thymus and bone marrow. Shown are representative FACS-plots of three experiments with similar outcome. **(e)** The mean fluorescence intensity (MFI) was calculated from one representative experiment out of two with identical outcome (*n*=4). **(f)** Flow cytometric measurement of ß-Gal activity in splenic CD8^+^ DCs. Ppef2^lacZ/lacZ^ reporter mice (red, uninjected) were injected intravenously with LPS either 4h (blue) or 12h (black) before analysis. Shown are FACS-plots and statistics, where the ß-Gal signal of reporter mice was subtracted from the ß-Gal background signal of control mice (*n*=18 mice). Statistical analyses were performed by using Student’s t-test, with *p<0.05; **p<0.01; ***p<0.001.

### Ppef2-deficient mice have normal numbers of DCs

We next analyzed if lack of Ppef2 would have an impact on the composition of the DC-compartment or other cells *in vivo*. However, we could not detect differences as compared to Ppef2^+/+^ mice, when we analyzed frequencies and cell numbers of DC subsets in spleens, lymph nodes or thymi of Ppef2^−/−^ mice (Fig. 3a). Also analysis of DC subsets in the lung, intestinal lamina propria, liver and skin dermis could not reveal statistically significant differences of Ppef2^−/−^ DCs subsets *in vivo* (Fig. Suppl. 2). Further analysis of other hematopoietic cell subsets (Suppl. Fig. 3) or MHCII^−^CD11 c^−^CD43^+^sirpa^+^ DC-precursors in bone marrow and spleen (Fig. 3b) also showed no differences between Ppef2^−/−^ and wild-type controls. The spleen architecture of Ppef2^−/−^ animals and the positioning of DCs was normal as revealed by histological immunofluorescence analyses (Fig. 3c). Taken together, lack of Ppef2 did neither alter the composition of DC subsets nor the haematopoietic compartment in general.

**Figure 3.**
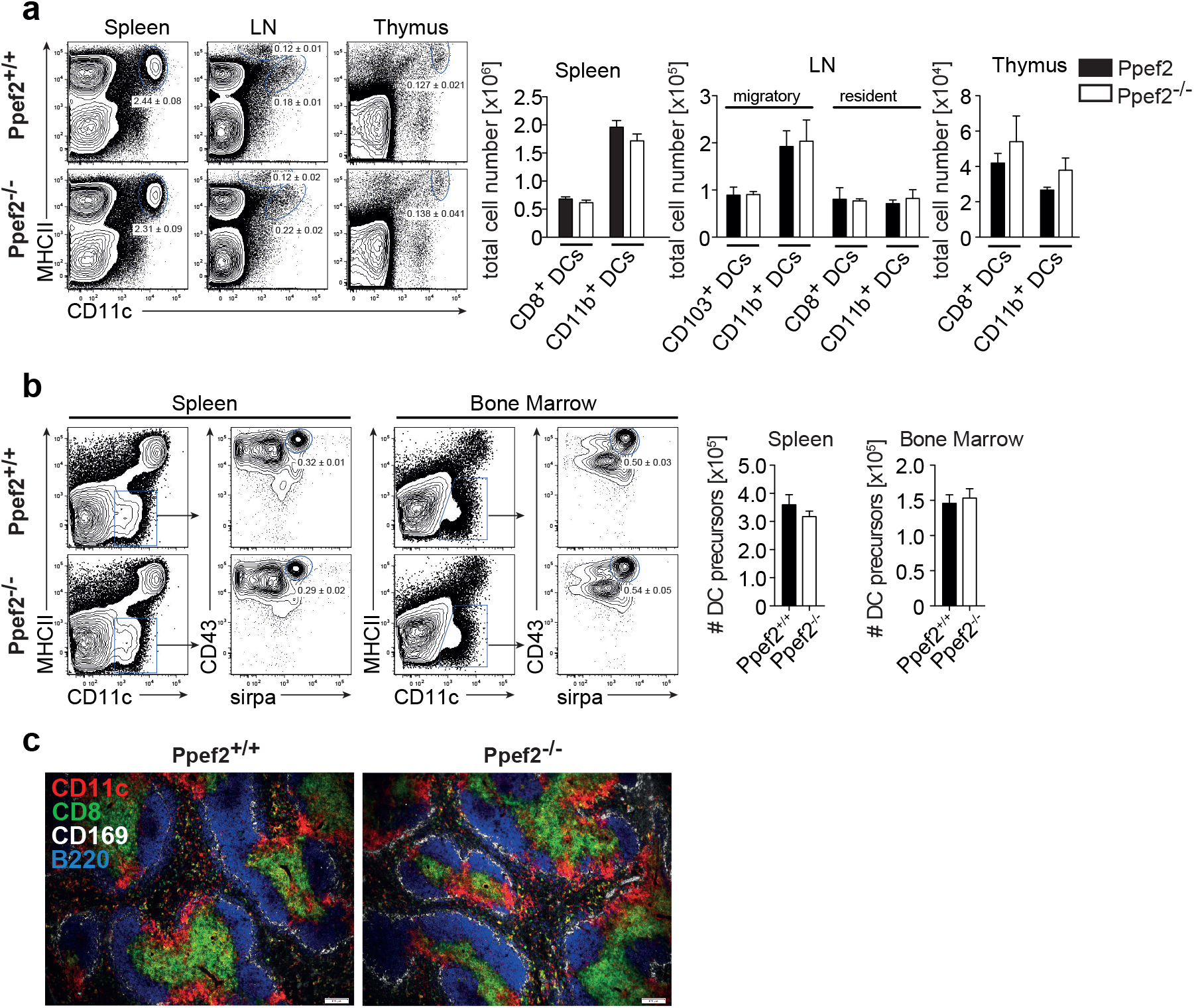
Ppef2^−/−^ -mice have normal frequencies of DCs and DC-precursors. **(a)** DCs of spleen, LN and thymus were stained with CD11c and MHCII. Shown are representative FACS plots with the average DC percentages ± SEM of pooled data (spleen, 8 experiments (*n*=36); sLN, 4 experiments (n=23); thymus, 2 experiments (*n*=6) as well as the corresponding total cell numbers. **b)** The frequency of DC precursors from spleen (*n*=3) and bone marrow (*n*=4). Shown is one representative experiment out of two with similar outcome. Statistical analysis of (**a**) and (**b**) was performed using Student’s t-test and all p-values were above 0.05. **(c)** Acetone fixed spleen sections of Ppef2^+/+^ and Ppef2^−/−^ mice were stained with antibodies against CD11c (red), CD8 (green), CD169 (white), and B220 (blue). Scale bars represent 100 μm.

### Lack of Ppef2 causes an increased rate of apoptosis and decreased survival rate in DCs

As Ppef2 has been linked functionally to cell survival and apoptosis in previous *in vitro* studies in human cells (Kutuzov, Bennett et al., 2010), we next tested if Ppef2^−/−^ DCs would show increased apoptosis. We analyzed CD11c^+^MHCII^+^ DCs from spleens for presence of cleaved caspase 3 and detected significantly more Casp3^+^CD11c^+^MHCII^+^ DCs in Ppef2^−/−^ mice as compared to wild type controls (Fig. 4a, b). More detailed analysis showed that Ppef2^−/−^ CD8^+^ DCs contained the highest frequencies of Casp3^+^ apoptotic cells, but also CD11b^+^ DCs were affected to some extent (Fig. 4a, b). In contrast, other lymphocytes from Ppef2^−/−^ mice such as B cells, T cells, macrophages or monocytes did not show any elevated levels of Casp3 (Fig. 4b). Taken together, these findings suggest that the absence of Ppef2 leads to increased rates of apoptosis in CD8^+^ DCs. We next wondered if this would influence their competitive behaviour *in vivo*. To test this, we chose a competitive situation and generated radiation bone marrow chimeras, which were reconstituted with a mix of bone marrow derived from CD45.1 ^+^ Ppef2^+/+^ and CD45.1^−^ Ppef2^−/−^ mice at a 1:1 ratio. The analysis of these chimeras showed that CD45.1^−^ Ppef2^−/−^ CD8^+^ DCs were non-competitive as compared to Ppef2^+/+^ CD8^+^ DCs in the same animal, as reconstitution was significantly less efficient for Ppef2^−/−^ CD8^+^ DCs as compared to Ppef2^+/+^ CD8^+^ DCs (Fig. 4c). This effect was specific for CD8^+^ DCs, as Ppef2^−/−^ CD11b^+^ DCs developed normally and generated frequencies comparable to those of Ppef2^+/+^ CD11b^+^ DCs (Fig. 4c). Therefore, although under non-competitive conditions Ppef2^−/−^ DCs developed in normal numbers (Fig. 3), they were outcompeted by Ppef2^+/+^ DCs during reconstitution upon lethal irradiation.

**Figure 4.**
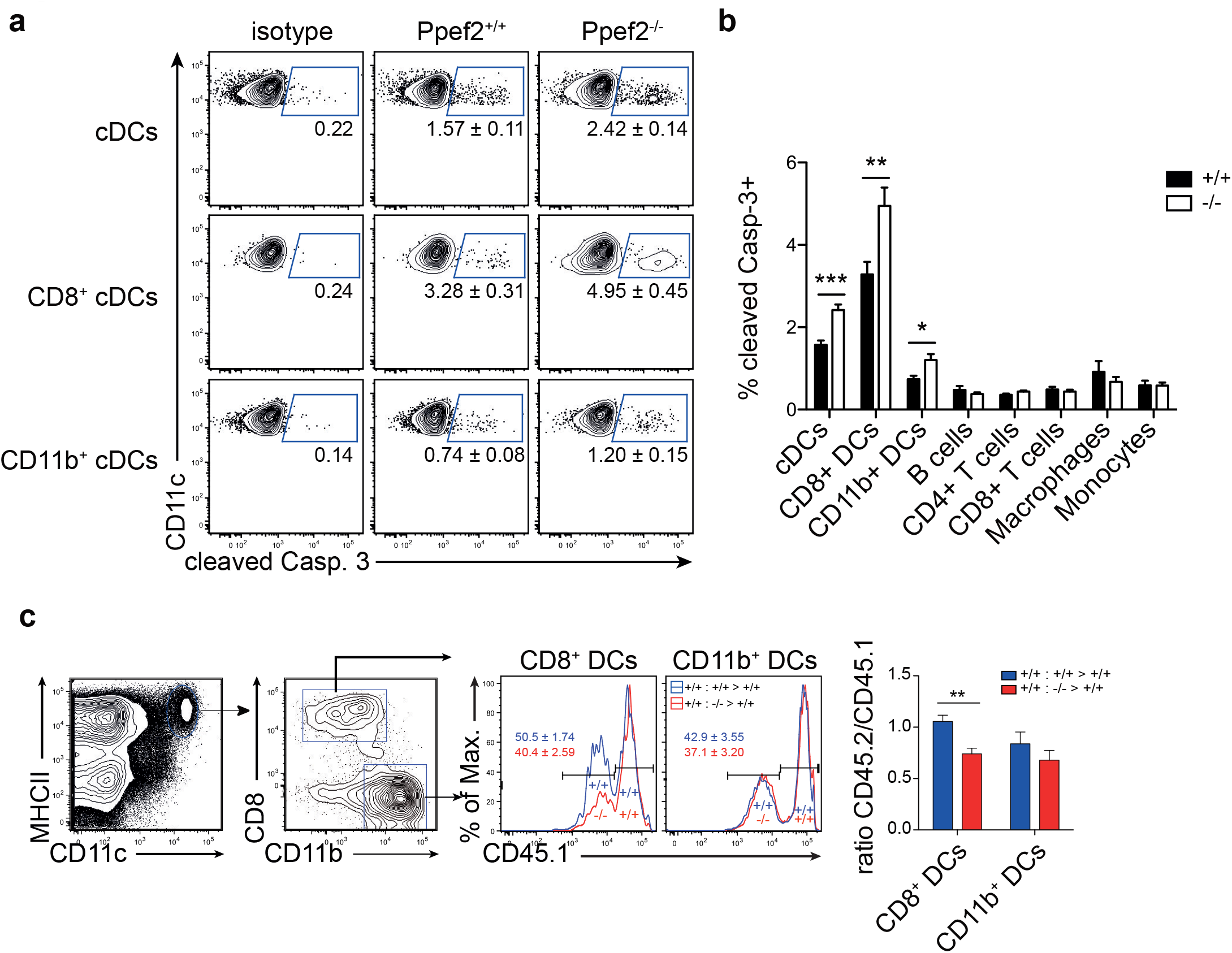
Ppef2-deficiency causes increased levels of cleaved caspase-3 in DCs. **(a)** Splenocytes were stained intracellularly for cleaved caspase-3 and analyzed by flow cytometry. Shown are representative FACS-plots of two experiments with similar outcome (*n*=4 each). The corresponding statistics of the pooled data (*n*=8) are shown in **(b)** together with the statistical analysis of other cell types. Statistical analysis was performed using Student’s t-test, with *p<0.05; **p<0.01; ***p<0.001. **(c)** Mixed bone marrow chimeras were produced by irradiation of CD45.1^+^ recipients and reconstitution with a 1:1 mix of Ppef2^+/+^ (CD45.1^+^) and Ppef2^+/+^ (CD45.2^+^) bone marrow (+/+ : +/+ > +/+), or a 1:1 mix of Ppef2^+/+^ (CD45.1^+^) and Ppef2^−/−^ (CD45.2^+^) bone marrow (+/+ : −/− > +/+). Mixed bone marrow chimeras were analyzed 8-10 weeks after reconstitution by gating on CD11c^+^MHCII^+^CD8^+^CD11b^−^ DCs, CD11c^+^MHCII^+^CD8^−^CD11b^+^ DCs and CD45.1. Shown are representative FACS-plots of three independently performed experiments with similar outcome (*n*=11). The ratios of CD45.2^+^ to CD45.1^+^ DCs were calculated from pooled data (*n*=11 mice) and statistical analysis was performed using Student’s t-test, with **p<0.01.

### Transcriptome analysis of Ppef2^-/-^ CD8^+^ DCs reveals regulation of apoptosis and cell survival genes

To analyse if absence of Ppef2 might alter gene expression which eventually controls DC survival, we performed mRNA sequencing of sorted spleen CD8^+^ DCs (Fig. 5a). A total of 10766 genes were detected and the read counts for Ppef2 were used as internal control. Ppef2^−/−^ DCs did not show any Ppef2-reads above background (Fig. 5a), confirming the efficiency of the Ppef2-knockout. Analysis of differentially expressed genes revealed 13 down- (Fig. 5a, blue) and 8 up-regulated genes (Fig. 5a, red) that were at least two-fold different with a p-value ≦ 0.01. Among the differentially regulated genes, *tripartite motif-containing 2* (Trim2) was downregulated more than 2-fold (Fig. 5b); this gene is of particular interest because it binds to the phosphorylated form of the pro-apoptotic Bim, mediating its ubiquitination for degradation (Thompson, Pearson et al., 2011). Delta-like ligand 4 (Dll4), a ligand for Notch (Shutter, Scully et al., 2000), was down-regulated 2.8-fold in Ppef^−/−^CD8^+^ DCs (Fig. 5b). Notch signalling is known to be crucial for lymphocyte development and function (Maillard, Fang et al., 2005) and signaling via Notch maintains CD11b^+^ DCs (Caton, Smith-Raska et al., 2007). To validate the mRNA sequencing results, we performed a qPCR of Trim2 and Dll4 on sorted CD8^+^ DCs of the spleen (Fig. 5c). Both mRNAs were down-regulated significantly in Ppef2^−/−^ CD8^+^ DCs (Fig. 5c), confirming the results from mRNA-sequencing (Fig. 5a, b).

**Figure 5.**
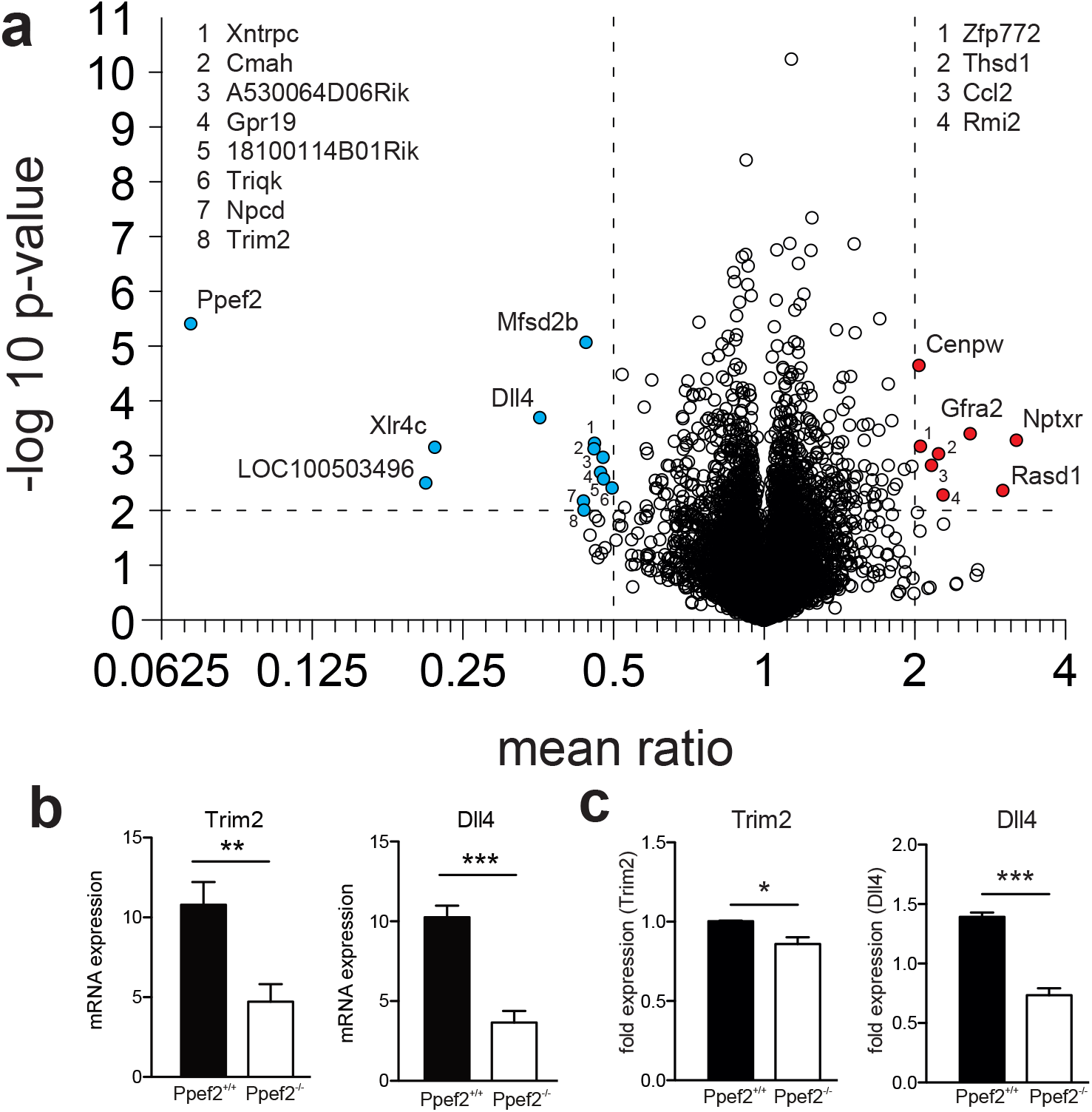
RNA-sequencing reveals changes in RNA-expression of Ppef2^−/−^ CD8^+^ DCs. **(a)** CD8^^+^^ DCs were purified by flow cytometry from cell suspensions of 3 pooled spleens as live MHCII^+^CD11c^+^CD11b^−^CD8^+^ cells to purity of >95%. 15 spleens from Ppef2^+/+^ or Ppef2^−/−^ mice were used to generate 5 samples each for RNA-sequencing. Shown is the volcano plot analysis of sorted CD8^+^ DCs. Fold change of -2 (**a**, blue) and ^+^2 (**a**, red), and a p-value ≤ 0.01 were chosen as cut-off. Ppef2, protein phosphatase EF-hands 2; LOC100503496, uncharacterized transcript LOC100503496; Xlr4c, X-linked lymphocyte-regulated 4C; Dll4, delta-like ligand 4; Trim2, tripartite motif-containing 2; Npcd, neuronal pentraxin chromo domain; Mfsd2b, major facilitator superfamily domain containing 2B; Cmah, cytidine monophospho-N-acetylneuraminic acid hydroxylase; Xntrpc, Xndc1-transient receptor potential cation channel, subfamily C, member 2; A530064D06Rik, Riken cDNA A530064D06 gene; 1810014B01 Rik, Riten cDNA 1810014B01 gene; Triqk, triple QxxK/R motif containing; Nptxr, neuronal pentraxin receptor; Rasd1, RAS, dexamethasone-induced 1; Gfra2, glial cell line derived neurotrophic factor family receptor alpha 2; Rmi2, RMI2, RecQ mediated genome instability 2; Thsd1, thrombospondin, type I, domain 1; Ccl2, chemokine (C-C motif) ligand 2; Zfp772, zinc finger protein 772; Cenpw, centromere protein W. **(b)** Boxplots represent normalized expression with 0,1 quantile, 0.9 quantile and all single points (each group *n*=5) **p<0.01, ***p<0.001 for Trim2 and Dll4 in Ppef2^+/+^ and Ppef2^−/−^ cells. **(c)** qPCR of Trim2 and Dll4 in sorted spleen CD8^+^ DCs. Spleen DCs were sorted as CD11c^+^MHCII^+^CD8^+^CD11b^−^ in three independent experiments and three Ppef2^+/+^ or Ppef2^−/−^ mice were pooled for every sort. Statistical analysis was performed using Student’s t-test, with *p<0.05; ***p<0.001.

### Ppef2^−/−^ DCs have elevated levels of cytoplasmic pro-apoptotic Bim

We next analyzed protein levels of Trim2 by Western blot analyses but could not detect significantly different expression between Ppef2^−/−^ and Ppef2^+/+^ BMDC (data not shown). Next we performed western blot analyses of Bim in GM-CSF cultured BMDCs (Fig. 6a) and sorted spleen CD8^+^ DCs (Fig. 6b). As Bim increases after stimulation with TLR ligands (Chen et al., 2007), we stimulated BMDCs with LPS as positive control. All three Bim isoforms, Bim_EL_, Bimi___, and Birn_β_ can promote apoptosis (O’Connor, Strasser et al., 1998) and could be readily detected (Fig. 6a, b). Quantification of Bim relative to the GAPDH loading controls showed that in unstimulated Ppef2^−/−^ BMDCs more Bim of all isoforms was present as compared to controls (Fig. 6a). Bim levels in unstimulated Ppef2^−/−^ BMDCs were similar to the increased levels found in LPS-induced Ppef2^+/+^ BMDCs (Fig. 6a). Stimulation of Ppef2^−/−^ DCs with LPS could increase Bim only marginally, while it increased in Ppef2^+/+^ wt DCs as described previously (Chen et al., 2007) (Fig. 6a).

**Figure 6.**
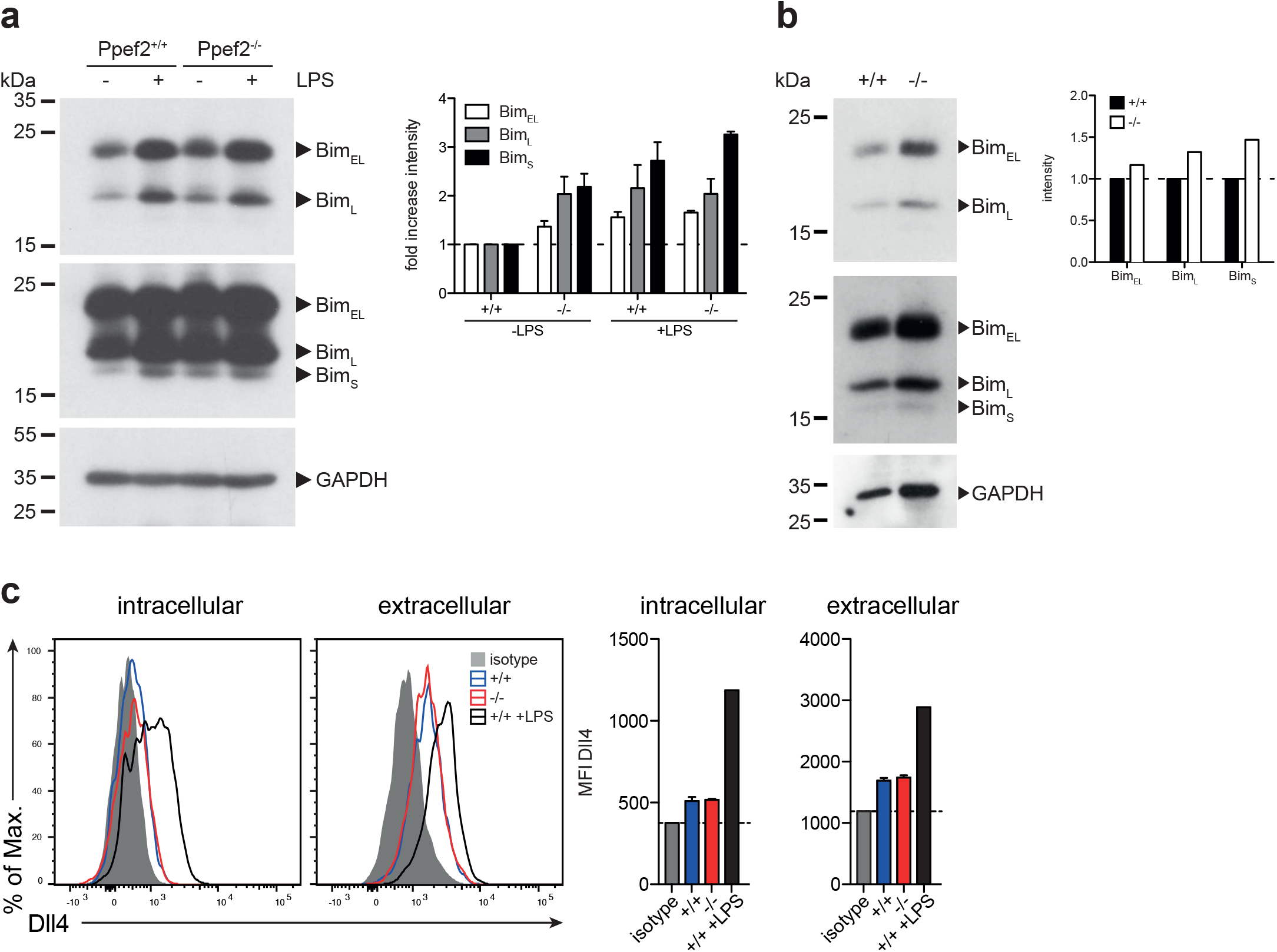
Ppef2^−/−^ DCs have elevated levels of pro-apoptotic BIM. **(a)** Western blot for Bim was performed with 30μg cell lysates of GM-CSF BMDCs either unstimulated or LPS matured. Exposure for 30s allowed detection of BimEL and BimL isoforms (**a** left, top panel); 5min exposure revealed the BimS isoform in addition (**a** left, middle panel). Intensities of the bands for all three Bim isoforms were quantified relative to the GAPDH loading control (**a** left, lower panel) by using ImageJ and fold increase was calculated relative to the untreated Ppef2^+/+^ control (**a**, right hand panel). Two western blots were performed and shown are pooled normalized intensities. **(b)** Western blot for Bim was performed with 30μg cell lysates of FACS-sorted CD8^+^ DCs of the spleens of Ppef2^+/+^ and Ppef2^−/−^ mice. Exposure of the membrane was performed like in (a). Intensities of the bands for all three Bim isoforms were quantified relative to the GAPDH loading control by using ImageJ and fold increase was calculated relative to the Ppef2^+/+^ control. Shown is the result of one experiment with pooled samples of three mice per group. **(c)** Surface staining of Dll4 was performed with fluorescently labelled antibody and protein abundance was measured by flow cytometry. Shown are representative FACS plots (left) and mean ^+/−^ SEM of the mean fluorescence intensity of *n*=3 mice.

In primary CD8^+^ DCs from spleen, protein abundance of all three Bim isoforms was increased in Ppef2^−/−^ CD8^+^ as compared to Ppef2^+/+^ CD8^+^ DCs, when quantified relative to the GAPDH loading controls (Fig. 6b). This suggests that Bim-mediated apoptosis might cause the alterations observed in Ppef2^−/−^ DCs and cause their competitive disadvantage (Fig. 4c).

Although expression of Dll4 is generally low in cDCs (Heng & Painter, 2008), we saw changes of Dll4-mRNA-expression in CD8^+^ cDCs of Ppef2^−/−^ mice (Fig. 5a, b). Therefore, we next analysed Dll4-protein levels. Spleen cDCs showed only minor intracellular staining over background (isotype) control, with slightly elevated surface expression levels (Fig. 6c). As published previously, LPS-treatment augmented DII4-expression significantly (Skokos & Nussenzweig, 2007). However, Ppef2^+/+^ and Ppef2^−/−^ DCs showed comparable Dll4 levels, indicating that the difference in Dll4-mRNA-expression could not be confirmed on surface protein levels. Taken together, Ppef2-deficiency seems to cause increased levels of pro-apoptotic Bim in CD8^+^ DCs.

### Ppef2^−/−^ DCs display decreased capacity of CD8^+^ T cell priming

The life-span of DCs directly influences antigen-specific expansion of T cells (.Chen et al., 2006, Nopora & Brocker, 2002). We therefore wondered whether Ppef2-deficiency would alter antigen-presentation and T cell priming *in vivo*. As infection models with pathogens or protein immunization with adjuvants contain TLR-signals which rapidly downregulate Ppef2 in Ppef2^+/+^ DCs (Fig. 1c, 2f), such models render Ppef2^−/−^ and Ppef2^+/+^ mice quite similar with respect to low Ppef2-levels. CD40-crosslinking did not induce Ppef2-downregulation (Fig. 1c). Therefore, we adoptively transferred TCR-transgenic CD90.1^+^ OT-I T cells into Ppef2^+/+^ or Ppef2^−/−^ mice and injected ovalbumin (OVA) as cognate antigen together with an agonistic anti-CD40 mAb for DC-maturation (Bonifaz, Bonnyay et al., 2002). Here, cross-presentation by CD8^+^ DCs should be the main mechanism for CTL-priming. While OT-I cells expanded strongly in the spleens of Ppef2^+/+^ mice, only marginal OT-I expansion was detected in Ppef2^−/−^ mice (Fig. 7a, upper panel). The lack of efficient OT-I cross-priming in Ppef2^−/−^ mice was evident from the percentage of OT-I cells found on day 7 post priming in the spleens of mice, as well as from the total OT-I numbers (Fig. 7a, upper panel). When OT-I cells were restimulated with the cognate OVA257 peptide *in vitro*, only very low frequencies and total numbers of CD107^+^IFN-γ^+^ OT-I effector T cells were found in spleens of Ppef2^−/−^ mice (Fig. 7a, lower panel). In contrast, OT-I cells primed in Ppef2^+/+^ mice were readily producing IFN-γ^+^ (Fig. 7a, lower panel). We next tested antigen presentation in absence of inflammatory stimuli and injected OVA in PBS. 3 days later Ppef2^−/−^ mice showed a significantly reduced OT-I T frequency and nearly 40% reduced OT-I T cell numbers (Fig. 7b). This data suggests that cross-presentation of OVA-protein by immature DCs in absence of inflammatory stimuli as well as by CD40-matured DCs is severely reduced in Ppef2^−/−^ mice.

**Figure 7.**
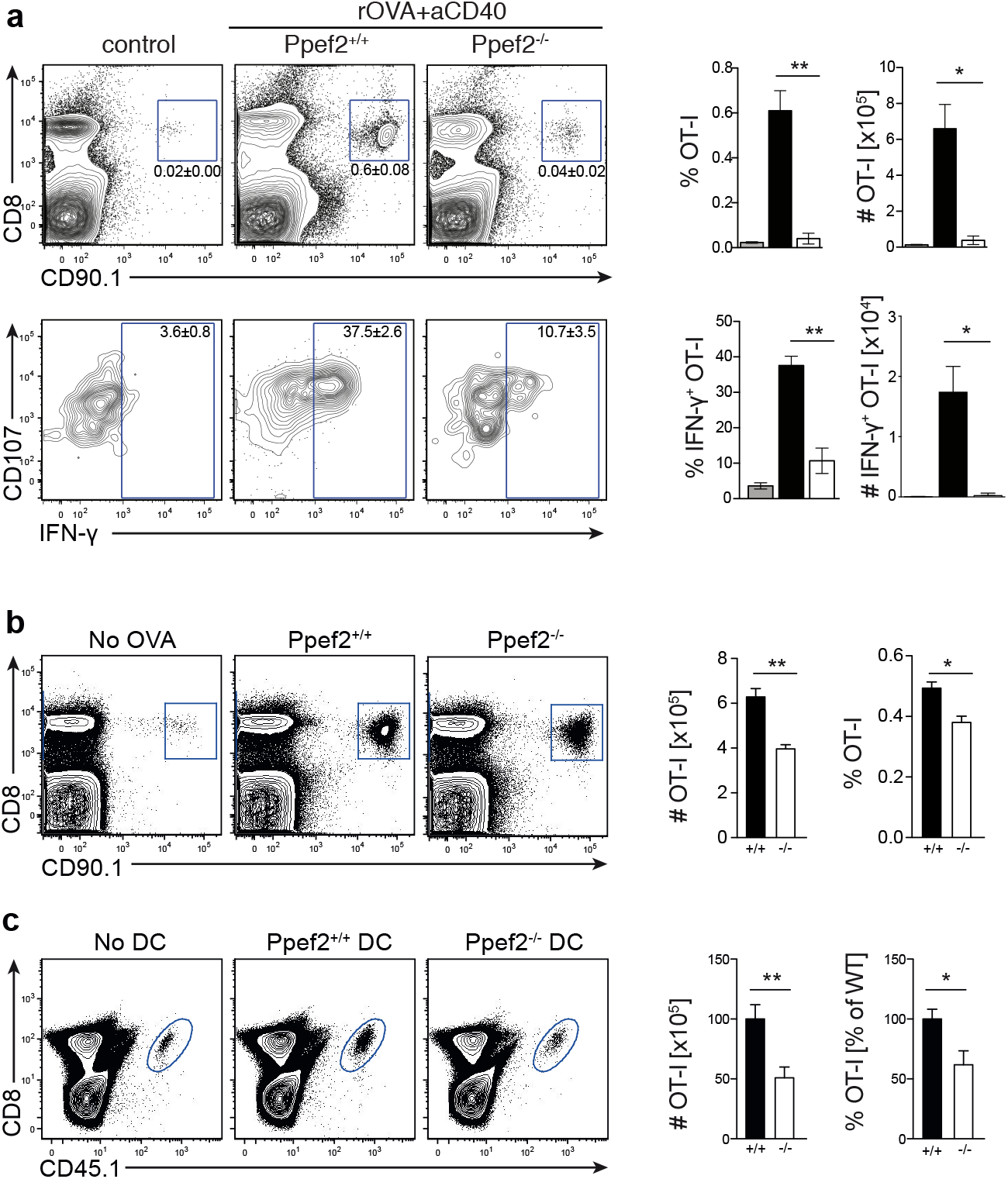
Ppef2^−/−^ DCs display decreased CD8^+^ T cell priming capacities. **(a)** 5x10^5^ congenitally marked CD90.1^+^ OT-I T cells were adoptively transferred either in Ppef2^+/+^ or Ppef2^−/−^ mice, which received 30μg OVA protein and 100μg CD40-specific mAb intravenously one day later. On day 7 post immunization spleens were analyzed for frequencies and total cell numbers of CD90.1^+^CD8^+^ OT-I cells as shown in the respective bar graphs (upper panel). Spleen cells were cultured *in vitro* with OVA257 peptide in the presence of CD107a-specific mAb and subsequently stained for CD8 and IFN-γ (lower panel) to determine IFN-γ-producing OT-I T cell frequencies and total cell numbers (bar graphs). Shown are results from one experiment out of two with similar results and *n*=4 mice per group. **(b)** 5x10^5^ congenitally marked (CD90.1) OT-I T cells were adoptively transferred either in Ppef2^+/+^ or Ppef2^−/−^ mice, which received 100μg OVA protein intravenously one day later. Mice were sacrificed three days after OVA administration and spleens were analyzed by flow cytometry for OT-I T cell proliferation. Shown are representative FACS-plots of OT-I T cells of one out of two independently performed experiments with similar outcome. Data from two independent experiments were pooled (*n*=6) to determine the total cell number and frequencies of CD90.1^+^CD8^+^ OT-I T cells. **(c)** 1.5x10^5^ Ppef2^+/+^ or Ppef2^−/−^ DCs were pulsed with OVA257 peptide and transferred i.v. into H-2K^bm1^ recipient mice. 24h later H-2K^bm1^ OT-I T cells were transferred and mice were analyzed 3 days later for presence of CD45.1^+^CD8^+^ OT-I T cells by flow cytometry. Shown are representative FACS-plots of the gated CD45.1^+^CD8^+^ OT-I T cells. Two independently performed experiments were pooled (*n*=6) to determine the frequency and total cell number of CD45.1^+^CD8^+^ OT-I T cells from the gates shown in (c). Statistical analysis was performed using Student’s t-test, with *p<0.05; **p<0.01.

To exclude differential antigen uptake or processing of OVA antigen by Ppef2^−/−^ DCs *in vivo*, we next transferred OVA257-peptide pulsed Ppef2^−/−^ or Ppef2^+/+^ DCs into H-2K^bm1^ recipient mice. Here, direct peptide loading of DCs avoids the necessity for antigen-processing, while usage of H-2K^bm1^ mice as recipients does not allow peptide presentation by cells of the host recipient mice, as the H-2K^bm1^ mutation of the H2-K^b^ gene leads to a failure to present OVA257 on MHC class I (Nikolic-Zugic & Carbone, 1990). Adoptively transferred CD45.1^+^H-2K^bm1^ OT-I T cells showed strongly reduced frequencies and total cell numbers when stimulated by Ppef2^−/−^ DCs as compared to Ppef2^+/+^ DCs (Fig. 7c). Taken together, our data suggests that Ppef2 expression in CD8^+^ DCs ultimately contributes to efficient cross-presentation of protein antigen.

## Discussion

Currently very little is known about Ppef2. In this study we show that among hematopoietic cells specifically CD8^+^ DCs express the phosphatase Ppef2, which is down-regulated upon DC-maturation by TLR-ligands, but not by CD40-engagement. Down-regulation of Ppef2 increases apoptosis and limits antigen-presenting cell functions of CD8^+^ DCs.

Transcriptional profiling showed that Ppef2 is expressed with high specificity in CD8^+^ DCs and confirmed the microarray data available from the Immgen consortium (Heng & Painter, 2008). Although CD11b^+^ DCs and monocytes were also reported to express very low levels of Ppef2 mRNA, this did not translate into protein as determined by β-galactosidase, which is included in the gene-trap cassette of Ppef2^−/−^ mice. However, CD103^+^ DCs, which are the tissue counterparts of lymph node and spleen CD8^+^ DCs (Bursch, Wang et al., 2007, del Rio, Rodriguez-Barbosa et al., 2007, Merad et al., 2013) and show some Ppef2-mRNA expression according to the Immgen database (Heng & Painter, 2008), do not express significant amounts of Ppef2 (ß-Gal)-protein. CD11b^+^ DCs of the thymus, but not of other tissues, are the only other Ppef2^+^ cell type in addition to CD8^+^ DC. As Ppef2 shares promoter organization with CD11c and the respective promoter motive is sufficient to drive DC-specific gene expression in all CD11c^+^ DCs (Edelmann et al., 2011), the CD8^+^ DC-specific Ppef2-expression is surprising. This suggests that Ppef2-expression is most likely controlled on additional levels leading to differential expression in distinct DC subpopulations. More specifically, CD8^+^ and CD103^+^ DC subsets differentiate from the same Ppef2-negative pre-DC precursor into different environments. While CD8^+^ DCs are mostly resident in lymphoid organs, CD103^+^DCs develop in barrier organs such as skin and mucosal tissues. Here, CD103^+^ DCs might routinely encounter TLR-signals from skin, lung and intestinal microbes, which could eventually cause down-regulation of Ppef2 in the steady-state. In contrast, CD8^+^ DCs in lymph nodes and spleen should not readily receive TLR-signals due to their localization and might therefore continue to express Ppef2.

In the case of an infection, exposure of CD8^+^ DCs to TLR-ligands also down-regulates Ppef2-expression in mature DCs. This is in contrast to CD40-signals; although both signalling pathways utilize TNF-associated factor (TRAF) 6, CD40 and TLR-signalling leading to NF-κB activation vary from each other (O’Sullivan & Thomas, 2002). CD40 mediated TRAF-signalling involves also TRAF2 and 3 (Tsukamoto, Kobayashi et al., 1999), while TLR-signalling involves IL-1R-associated kinases (IRAKs). In addition, CD40 signalling can result in protein kinase B (Akt) activation (Arron, Vologodskaia et al., 2001), which may promote survival, while TLR signalling was shown to induce JUN N-terminal kinase (JNK), which may cause apoptosis (Hull, McLean et al., 2002). This data fits to the hypothesis that CD40 induces pro-survival signalling, whereas TLR signaling is rather pro-apoptotic and involves mitogen-activated protein kinases, which regulate Ppef2 expression and possibly promote apoptosis. In contrast, pro-survival TRAF6-CD40 signalling might utilise Akt not altering the expressing of Ppef2. We observed down-regulation of Ppef2 expression also after spontaneous maturation of DCs in culture and other studies have shown similarities between LPS activated and spontaneously activated cells *in vitro*, when analysing microarray data (Riepsaame, van Oudenaren et al., 2013). The majority of changes in gene expression during DC-maturation are reductions of expression (Sanchez, McWilliams et al., 2007). Therefore, it is difficult to conclude whether loss of expression of Ppef2 in mature DCs is simply a consequence of co-downregulation of other genes, which may be dispensable for mature DCs, or whether it is a specific regulation in order to limit the lifespan of DCs.

The only functional Ppef2-study published so far investigated the anti-apoptotic role of Ppef2 in oxidative stress by suppression of Apoptosis signal-regulating kinase 1 (Ask1) in immortalized cell lines (Kutuzov et al., 2010). Ppef2 caused potent dephosphorylation of Ask1 at the threonine residue 838, suppressing Ask1 activity (Morita, Saitoh et al., 2001). When Ask1 and Ppef2 were co-expressed *in vitro*, suppression of Ask1 by Ppef2 led to a decrease in caspase-3 activation (Kutuzov et al., 2010). It was concluded that the phosphatase Ppef2 functions to control Ask1 activity to prevent apoptosis under oxidative stress conditions. Ppef2 is also up-regulated in mouse hippocampus after hypoxia-ischemia, along with several other transcripts associated with neuroprotection (Gilbert, Costain et al., 2003). However, western blot analyses of Ppef2^−/−^ DCs revealed no changes in Ask1 expression or phosphorylation (data not shown). Instead, mRNA-sequencing revealed Trim2 and Dll4 being expressed at reduced levels in Ppef2^−/−^ CD8 DCs.

Trim2, an E3 Ligase, ubiquitinates pro-apoptotic Bim for proteasomal degradation (Thompson et al., 2011). Ppef2^−/−^ DCs showed an increase in Bim, for which a decrease in Trim2 could be responsible, although we were unable to confirm alterations of Trim2 protein levels in Ppef2^−/−^ DCs (data not shown). In conclusion, we showed that loss of Ppef2 led to a decrease of Trim2 mRNA which can influence the abundance of Bim. Being a regulator of DC apoptosis, Bim would therefore be responsible for the increase in apoptotic active caspase-3^+^ DCs observed in Ppef2^−/−^ mice.

Interestingly, we also observed increased caspase-3 levels in CD11b^+^ DCs of Ppef2^−/−^ mice *in vivo*, although these DCs do not express Ppef2. Eventually indirect secondary effects or the uptake of apoptotic cleaved caspase-3^+^CD8^+^ DCs by CD11b^+^ DCs might account for this finding. Recently, it has been shown that oxysterol-metabolizing enzyme expressing CD8^+^ (XCR1 ^+^) DCs do influence homeostasis and positioning of CD11 b^+^ (DCIR2^+^) DCs (Lu, Dang et al., 2017), indicating a possible cross-talk between both DC-subsets. However, the exclusivity of the competitive disadvantage observed for CD8^+^, but not CD11 b^+^ DCs, during bone marrow chimera reconstitution suggests that Ppef2-deficiency directly affects CD8^+^ DCs only. Although the rate of apoptosis was increased in CD8^+^ DCs from Ppef2^−/−^ mice, we did not find diminished total numbers or frequencies of CD8^+^ DCs. Also short-term or pulse-chase BrdU-labelling experiments could not reveal statistically significant differences for DC proliferation or turnover, which might compensate for increased rates of DC-death (data not shown). As only 5% of all DCs proliferate in steady-state (Kabashima et al., 2005), BrdU-incorporation might not be sensitive enough to reveal subtle changes in compensatory proliferative rates. Also serum levels of Flt3L were not elevated, as reported in mice which lack DCs due to expression of diphteria toxin (Birnberg, Bar-On et al., 2008). Eventually, alterations in precursor-product relations might account for compensatory effects. However, we did not detect elevated numbers of pre-DCs, nor did they show an increased proliferative rate (data not shown). Increased apoptosis of DCs should lead to decreased T cell priming capacity. In our experiments we avoided to activate DCs with TLR-ligands, as fully matured DCs naturally down regulate Ppef2 and as such would not be different from Ppef2^−/−^ DCs. T cell proliferation was substantially reduced upon priming by Ppef2^−/−^ DCs. Also cross-presentation of OVA-protein by CD40-matured CD8^+^ DC was strongly reduced in Ppef2^−/−^ mice, indicating that DCs have limited T cell priming ability when they lack Ppef2. This might hint for a DC-intrinsic mechanism, which could limit antigen-specific T cell responses for protection from excessive immune activation to avoid collateral tissue damage and autoimmunity, which has been observed with apoptosis-resistant DCs (Chen et al., 2007, Chen et al., 2006, Nopora & Brocker, 2002, Park et al., 2006, Stranges et al., 2007).

Although its sequence is highly conserved in humans, hPpef2 does not show a similar DC-specific expression pattern (Lattin, Schroder et al., 2008), but is rather restricted to the retina. Yet, Ppef2 was identified as a survival phosphatase also in human HELA-cells in an RNA interference screening (MacKeigan, Murphy et al., 2005). Eventually, other members of the PPP phosphatase family such as PPP3CC, which is expressed in human DCs (Su, Wiltshire et al., 2004), might play corresponding roles.

Taken together, the Ppef2-locus might be an attractive tool due to its specificity for CD8^+^ DCs and could help to work out developmental and functional differences between CD8^+^ vs. CD103^+^ DCs.

## Materials and Methods

### Mouse strains

Alleles targeting Ppef2 were produced for the EUCOMM and EUCOMMTools projects by the Wellcome Trust Sanger Institute. JM8A3.N1 embryonic stem cells targeting the Ppef2 locus were purchased at EUCOMM and microinjection in C57BL/6 oocytes was performed by the Transgenic Core Facility at the Max-Planck-Institute for Molecular Cell Biology and Genetics in Dresden, Germany. Chimerism was determined by coat color, verified by genotyping and chimeric mice were further crossed with C57BL/6 mice. Mice were analyzed in sex and age-matched groups with 8–13 weeks of age. Animal experiment permission was granted by animal ethics committee Regierung von Oberbayern, Munich, Germany. Ppef2^−/−^, Ppef2^+/+^ and OT-I mice were bred and maintained at the animal facility of the Institute for Immunology, Ludwig-Maximillians-Universität München. H2-K^bm1^ mice were bred and kept in the Institut für Medizinische Mikrobiologie, Immunologie und Hygiene at the TU Munich.

### *In vitro* cell cultures

For GM-CSF BMDC cultures 10^7^ cells were plated in 10ml of GM-CSF containing medium (20 ng/ml GM-CSF). At day 3 of the culture, cells were harvested with Trypsin and again plated at a density of 5x10^6^ cells in GM-CSF medium. For analysis, cells were harvested at day 8 of the culture with cold PBS. For Flt3L cultures 3x10^6^ bone marrow cells were cultured with Flt3L medium and harvested at day 8 for analysis. Mature BMDCs were obtained by stimulating overnight with 2 μg/ml lipopolysaccharide (LPS, Sigma-Aldrich), 1 μg/ml Flagellin, 2.5 μg/ml Poly I:C, 1 μg/ml Pam3CSK4, 2.5 μg/ml CLO97, or 100 μg/ml anti-CD40, respectively.

### Bone marrow chimeras

To generate bone marrow-chimeras recipient mice were irradiated with two split doses of 550 rad using a Cesium source (Gammacell 40, AECl,Mississauga, Canada). Irradiated animals were reconstituted with 5x10^6^ BM cells, 1:1 mixed from CD45.1 ^+^ and CD45.2^+^ BM. To prevent infection, animals received 1.2g/l neomycin in water ad libitum for 4 weeks. Animals were analyzed 8–10 weeks after reconstitution.

### Flow cytometry analysis

Where possible, 2x10^6^ cells were used for every staining with titered antibodies in PBS containing 2% FCS and 0.01% NaN3 (fluorescence-activated cell sorting (FACS) buffer) and staining was carried out for 20 min at 4°C in the dark. Cells were washed once and used for direct acquisition on BD FACSCanto II. Dead cells were excluded using Aqua LIVE/DEAD Fixable Aqua DeadCell Stain Kit (Invitrogen, TermoFischer, Cat: L34957) or Zombie Aqua Fixable Viability Kit (BioLegend, Cat: 423102).

For the staining of cleaved caspase-3 cells were washed once and then resuspended in 200 μl 4% PFA for 15 min at room temperature in the dark. After washing twice with 1x fixation/permeabilisation buffer (BD) cells were blocked for 10 min with an anti CD16/32 antibody (Fc block, clone 2.4G2, BD) and 0.1% goat serum at 4°C. After 10 min of blocking, 100 μl anti cleaved caspase-3 antibody (clone D3E9, Cell Signaling) was added and cells were incubated for 30 min at 4°C. Unbound antibody was washed away and cells were incubated with 100 μl fluorochrome conjugated secondary antibody (goat anti-rabbit, Life Technologies) at 4°C for 30 min. Afterwards, cells were washed once and acquired at the FACS.

### Measurement of β-galactosidase (lacZ) expression and activity

Measurement of the expression and activity of the bacterial β-galactosidase gene (lacZ) was carried out using the FluoReporter lacZ Flow Cytometry Kit (Thermo Fisher Scientific Inc.) according to the manufacturers protocol. Therefore, the activity of the enzyme was measured by adding Fluorescein isothiocyanate (FITC) coupled substrate (fluorescein di-V-galactoside (FDG)) for 2 min at 37°C and stopping of the reaction by addition of cold staining medium. Followed by normal surface antibody staining, samples were analyzed using a FACSCanto II.

### Cell sorting

Cell sorting was either performed by staining the samples with fluorescently labelled antibodies and FACS-sorting the cells at a FACSAria or FACSFusion, or by staining the samples with magnetically labelled antibodies with MACS columns.

### T cell priming assays

0.5x10^6^ purified OT-I cells were transferred intravenously into congenic mismatched Ppef2^+/+^ and Ppef2^−/−^ mice that received 100μg grade IV OVA protein one day later i.v. *In vivo* T cell proliferation was analyzed three days after OVA injection staining for CD8 and CD90.1 or CD45.1, respectively. For immunization with anti-CD40 mAb, 100μg FGK4.5 (BioXCell, West Lebanon, USA) were co-injected together with 30μg OVA i.v.

For DC vaccinations, DCs were isolated from spleens of Ppef2^+/+^ and Ppef2^−/−^ mice as previously described (Mempel, Henrickson et al., 2004, Prlic, Hernandez-Hoyos et al., 2006). CD11c+ GFP^high^ splenocytes were sorted on a MoFlo™ XDP cell sorter, using fluorochrome-labelled antibodies specific for CD3 (500A2), CD19 (1D3), CD11c (N418), and propidium iodide (PI, Invitrogen) for live-dead discrimination. Sorted DCs were then pulsed with 1μg/ml OVA257 (SIINFEKL) peptide for 60min (37°C) and washed three times with an excess volume of PBS. 1.5x10^5^ OVA257 pulsed DCs were injected i.v. and transferred into H-2K^bm1^ recipient mice intravenously. 24 hours later 10^5^ H-2K^bm1+/+^ CD45.1+ naïve CD8+CD44^lo^ OT-I cells were transferred and T cell expansion was measured three days later.

### RNA-Isolation

Spleen CD11c^+^MHCII^+^CD8^+^CD11b^−^ DCs were sorted on a FACSAria five independent times with three pooled Ppef2^+/+^ or Ppef2^−/−^ mice per sort. For RNA isolation 5 × 10^5^ - 6 x 10^6^ cells per sample were harvested, subsequently lysed in TRIzol (Invitrogen), and total RNA was extracted with the miRNeasy kit (Qiagen) according to the manufacturers protocol. The quality of the RNA was assessed by measuring the ratio of absorbance at 260 nm and 280 nm using a Nanodrop 2000 Spectrometer (Thermo Scientific) as well as by visualization of the integrity of the 28S and 18S bands via a RNA analysis ScreenTape assay on a 2200 TapeStation instrument (Agilent).

### Western Blot analysis

Proteins were separated by SDS-PAGE (15%). Cell lysates were quantified using the Quant-iT Protein Assay kit (Molecular Probes). Equal amounts of protein were loaded onto 15% SDS gels, and GAPDH (clone 14C10, Cell signaling) was used as a loading control. After transferring to a nitrocellulose membrane, proteins were incubated with primary antibodies against Bim (clone C43C5, Cell Signaling) or GAPDH, washed, and incubated with secondary HRP-labelled antibodies (Jackson Immuno-Research Laboratories). Then, membranes were visualized using luminescent substrate ECL (GE Healthcare). Western blot bands were quantified using ImageJ software (National Institutes of Health).

### Statistics

For absolute cell numbers the percentage of living cells of a given subset was multiplied by the number of living cells as determined by CASY Counter. If not mentioned otherwise, significance was determined using the Student’s t-test and defined as follows: *P<0.05, **P<0.01 and ***P<0.001. Bar graphs show mean±s.e.m. for the group sizes as indicated in the figure legends.

### Data availability

Sequence data that support the findings of this study have been deposited with the primary accession code GSE98235 at https://www.ncbi.nlm.nih.gov/geo/query/acc.cgi?acc=GSE98235. The other data that support the findings of this study are available from the corresponding author upon request.

Supplemental Information includes additional Supplemental Experimental Procedures.

## Acknowledgements

This work was supported by the Deutsche Forschungsgemeinschaft SFB 914 A06 to S.S. and T.B., SFB1054 B03 to T.B. and SFB1054 B09 to D.B.; We thank Hervé Luche (CIPHE) for providing us with mouse FACS-data and analyses. J.L.S. is a member of the excellence cluster ImmunoSensation. We acknowledge the Core Facility Flow Cytometry at the Biomedical Center, Ludwig-Maxmilians-Universität München, for providing equipment and expertise.

## Author contributions

M.Z., T.U., X.C., C.R., C.B., L.G., C.S. conducted experiments; J.L.S. performed sequencing analysis; D.B. and V.B. provided adoptive transfer models and designed experiments; T.B. and S.S. designed experiments and wrote the paper.

